# A hierarchical framework for segmenting movement paths

**DOI:** 10.1101/819763

**Authors:** Wayne M. Getz

## Abstract

Comparative applications of animal movement path analyses are hampered by the lack of a comprehensive framework for linking structures and processes conceived at different spatio-temporal scales. Although many analyses begin by generating step-length (SL) and turning-angle (TA) distributions from relocation time-series data—some of which are linked to ecological, landscape, and environmental covariates—the frequency at which these data are collected may vary from sub-seconds to several hours, or longer. The kinds of questions that may be asked of these data, however, are very much scale-dependent. It thus behooves us to clarify how the scale at which SL and TA data are collected and relate to one another, as well as the kinds of ecological questions that can be asked. Difficulties arise because the information contained in SL and TA time series is not semantically aligned with the physiological, ecological, and sociological factors that influence the structure of movement paths. I address these difficulties by classifying movement types at salient temporal scales using two different kinds of vocabularies. The first is the language derived from behavioral and ecological concepts. The second is the language derived from mathematically formulated stochastic walks. The primary tools for analyzing these walks are fitting time-series and stochastic-process models to SL and TA statistics (means, variances, correlations, individual-state and local environmental covariates), while paying attention to movement patterns that emerge at various spatial scales. The purpose of this paper is to lay out a more coherent, hierarchical, scale-dependent, appropriate-complexity framework for conceptualizing path segments at different spatio-temporal scales and propose a method for extracting a simulation model, referred to as M^3^, from these data when at a relatively high frequencies (ideally minute-by-minute). Additionally, this framework is designed to bridge biological and mathematical movement ecology concepts; thereby stimulating the development of conceptually-rooted methods that facilitates the formulation of our M^3^ model for simulating theoretical and analyzing empirical data, which can then be used to test hypothesis regarding mechanisms driving animal movement and make predications of animal movement responses to management and global change.

## Introduction

The movement of organisms over landscapes can be usefully analyzed from various viewpoints. Four are suggested by the conceptual framework for movement ecology proposed by Nathan et al. (2008), depending on which questions constitute the focal interest—why move? how is movement executed? where to move? and how does the immediate environment, as well as the internal state of individuals, impact movement? Several more viewpoints arise when we focus on the structure of the movement path itself and how this structure may fill space at different spatio-temporal and, hence, ecological relevant scales (Levin 1992)—from the sequencing of various high-resolution movement elements to the emergence of seasonal home ranges and habitat utilization distributions (Worton 1987, Morellet et al. 2013, Torney et al. 2018, Wittemyer et al. 2019). The current lack of a comprehensive framework for linking structures and processes conceived at different spatio-temporal scales, however, hampers the synthesis of these viewpoints and of a comparative meta-analyses of similar questions addressed across disparate species and spatio-temporal scales. With this in mind, a more coherent, hierarchical, scale-dependent, appropriate-complexity framework than currently exists is laid out in this paper for conceptualizing path segments at different spatio-temporal scales and used to develop a model that captures the multimodal, correlated, random walk aspects present in relatively high frequency (i.e., minute-by-minute) relocation data. Of considerable importance is that this framework underscores the need to collect high resolution relocation data (minute or subminute sampling intervals) whenever possible.

Nathan et al. (2008) argued that movement arises from the interactions of several processes and motivations that can be understood through an analysis of external factors or stimuli, the internal state of an organism (e.g., see Morelle et al. 2015), and the organism’s mechanical and navigational capacities. These interactions then result in a movement process that may be characterized by a time series of relocation points that define movement paths (Siniff & Jessen 1969, Edelhoff et al. 2016). From these relocation time series, sequences of step lengths (SL) and turning angles (TA) can be extracted (Kareiva & Shigesada 1983, Turchin 1998) and organized into time-stamped pairs of values. Other data, such as accelerations (Nathan et al. 2012), audio (Sapir et al. 2010, Hurme et al. 2019, Northrup et al. 2019) and magnetometer (Williams et al. 2017, Chakravarty et al. 2019) data, as well as physiological measurements (e.g., temperature, blood glucose levels; Wilmers et al. 2015), may be recorded and used in the context of relocation point covariates. Additional covariates may be environmental measurements associated with resistance to movement caused by landscape topology and vegetation structure (Shepard et al. 2013, Getz & Saltz 2008, Spiegel et al. 2015), as well as local environmental features inferred from remotely sensed electromagnetic spectral data (e.g., visual and infra-red satellite imagery or lidar; Pettorelli et al. 2011, 2014, Bork & Su 2007).

In this paper, the question is addressed of how we may reconcile the fact that SL and TA distributions are not directly interpretable in terms of behavioral and ecological processes (e.g., resting, stalking, hiding, gathering); but, depending, on the spatio-temporal scale of the relocation data, phenomenological interpretations are made using scale-appropriate activity designators (Johnson et al. 2002, Owen-Smith & Goodall 2019, Giotto et al. 2015). A deeper understanding of the connections between path SL and TA time series and the designation of the so-called behavioral state of an individual at each of its relocation points requires that we find a way to reconcile narratives constructed using two different kinds of vocabularies. The first is the language of biological designators arising from behavioral and ecological concepts. Our primary tools here are the application of path segmentation methods (Nams 2014, Edelhoff et al. 2016, Seidel et al. 2018) that include hidden Markov models (HMMs; Michelot et al. 2016, Franke et al. 2004, Langrock et al. 2012, Zucchini et al. 2016) and behavioral change-point analyses (BCPA; Gurarie et al. 2009, 2016, Owen-Smith & Martin 2015). The second is the language of mathematical stochastic walks (Kareiva & Shigesada 1983, Bartumeus et al. 2005, Patterson et al. 2008) that treats movement relocation data as a time series of values and applies existing time-series methods (Chatfield 2016) to finding the parameter values that best fit selected time-series models to movement paths, with the possible inclusion of individual and environmental covariates (Jonsen et al. 2003, Preisler et al. 2004, Johnson et al. 2008, Fleming et al. 2014, Calabrese et al. 2016).

Moving between stochastic-walk formulations and biological designators in describing segments of movement paths is akin to translating between phonologic (e.g., Latin, cyrillic) and logographic (e.g., hieroglyphics, Chinese) scripts: much context-dependent interpretation needs to take place. But, in fact, the situation is much more dire, since we do not have access to every character—i.e., to the fundamental movement element (FuMEs, see glossary; Getz & Saltz 2008) of a movement path sequence. Rather, we only have access to some kind of “average” value across several characters or potential point-source values. The primary goal of this paper is to flesh out the temporal structure of biological designators of movement-path segments, provide a more coherent insights into the relationship between biological designators and stochastic-process formulations at different temporal scales, and provide a way for extracting useful stochastic process models from a hierarchically framed understanding of move path structure.

## A Pragmatic View of Path Segmentation

The view that large segments of individual movement paths can be parsed into strings of fundamental movement elements (FuMEs—see glossary; Fig. 1, Table 1) with characteristic frequencies and patterns of occurrence is highly idealized. Identifying FuMES is generally a biomechanical problem that is beyond the scope and capabilities of analyses based on relocation data alone. Other kinds of data, such as audio recordings, have been used to identify FuMEs: e.g., wingflaps of small birds (Sapir et al. 2010, Northrup et al. 2019) or footfalls of foraging ungulates (Northrup et al. 2019). If the frequency of our relocation data, however, spans strings of around 10-100 FuMEs, occurring either repetitively or in correlated groups of a couple of FuME types, then we may be able to reliably identify different strings of predominantly one type of FuME, or stereotypical mixed sequences of particular FuMES, which we call “metaFuMEs.” With this definition, for relatively high frequency relocation data, all the different metaFuMEs, in a set of metaFuME types, have the same fixed length equal to the time between consecutive relocation points and each type has its own characteristic bivariate SL and TA distribution, and auto- and cross-correlation time-series coefficients (Chatfield 2016). Such sets of metaFuMEs can then be used as a basis for representing different types of identifiable activities (e.g., foraging or focused travelling to a selected location), referred to as canonical activity modes (CAMs—see glossary; Getz & Saltz 2008).

**Table 1:** Hierarchical organization of a typical vertebrate life time track (LiTT). (Spatio-temporal scaling applies best to typical medium to large vertebrates, but not as well to small vertebrates, birds, or invertebrates.)

| Scale/<br>Segment | Time | Space | Categories | Data | Some approaches<br>and methods |
| --- | --- | --- | --- | --- | --- |
| *Constraint level: type of analysis* |  |  |  |  |  |
| **FuMEs\*** | 0.1–few<br>secs<br>*highly constrained: biomechanical analyses* | 0–1000<br>cms | stride, trot, run<br>twist, jump, flap | accelerometer<br>video | machine learning<sup>*†</sup>,<br>Newtonian<br>mechanics |
| meta-<br>**FuMEs\*** | resolution of<br>relocation data<sup>*‡</sup><br>*base level: movement ecology analyses* | average<br>step length | bivariate SL & TA<sup>¶</sup><br>distributions | SL & TA<br>means, variances<br>& correlations<sup>††</sup> | cluster analysis<br>ensemble filtering |
| short<br>duration<br>**CAMs†** | 0.1–10<br>mins<br>*highly flexible: sub-hourly/hourly analysis* | a few meters<br>to a few kms | resting, browsing,<br>riding thermals | secs to mins,<br>metaFuMEs &<br>covariate data | path<br>segmentation** |
| long<br>duration<br>**CAMs†** | 10–100<br>mins<br>*highly flexible: sub-diel analysis* | a few meters<br>to many kms | feeding<br>traveling | mins to hours,<br>short CAMs &<br>covariate data |  |
| **DARs‡** | fixed<br>24-hr<br>*time constrained: diel cycle analysis* | diel<br>range | central-place foraging<br>ranging | 24 hrs of data<br>CAM<br>sequences | periodograms<br>(Fourier, wavelets) |
| **LiMPs§** | several days to<br>a few months<br>*strongly constrained: lunar and seasonal cycle analyses* | home ranges<br>and their shifts<br>over time | philopatry, dispersal,<br>migration,<br>nomadic, recursion | DAR<br>sequences | clustering methods,<br>ideal-free/despotic<br>distribution<br>analyses |
| partial<br>or<br>whole<br>**LiTTs#** | paths<br>encompassing<br>several LiMPs<br>*moderately constrained: question-at-hand analysis* | habitat<br>types | territorial<br>ranging, migratory<br>nomadic | LiMP<br>sequences | GIS toolbox,<br>deep-learning<sup>*†</sup>,<br>pattern anal. |

\*Fundamental Movement Elements; †Canonical Activity Modes; ‡Diel Activity Routines
§Life-history Movement Phases; #Lifetime Tracks; ¶SL=step-length & TA=turning-angle
\*†Machine/deep learning methods can be applied at any scale but may be particularly useful here
\*‡Around 5 secs to 1 min
\*\*Includes Hidden Markov Models (HMMs) and Behavioral Change-Point Analyses (BCPA)
††See Section on **Stochastic Walk Statistics** in main text

**Figure 1:**
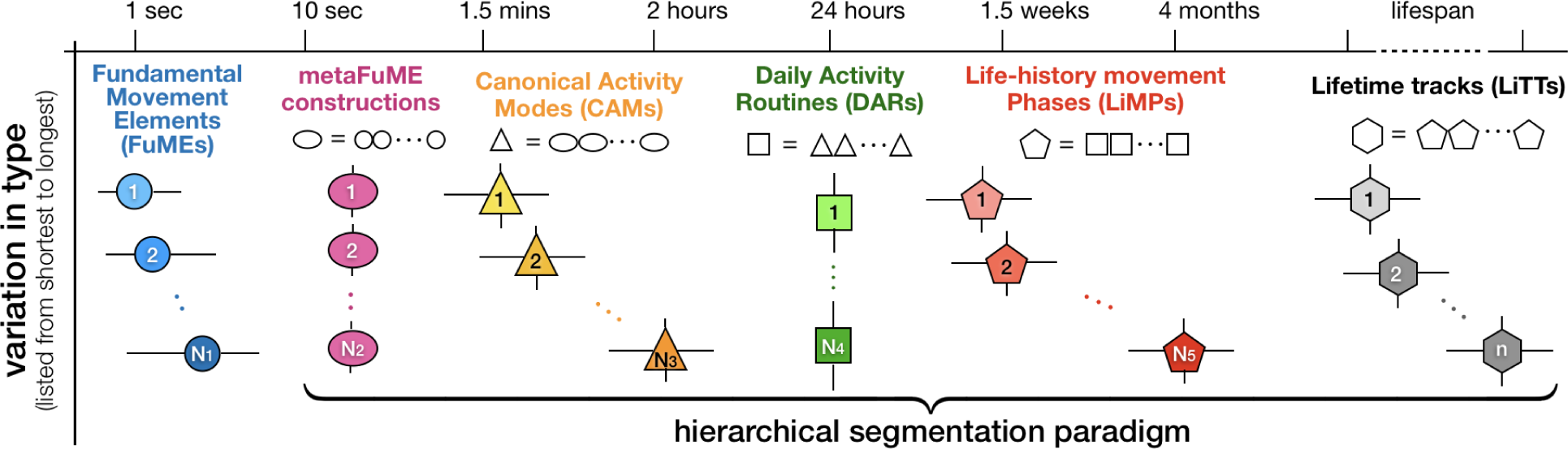
Scale-dependent path segmentation categories of lifetime track segments. The time scale applies to typical medium and large terrestrial animals and needs to be adjusted for aerial and marine species, and small animals, including invertebrates. Round (blue), oval (lilac), triangular (yellow to orange), square (green), pentagonal (red), and hexagonal (grey) icons are respectively used to represent *N*_1_ FuME types, *N*_2_ metaFuME types, *N*_3_ CAM types, *N*_4_ DAR types, *N*_5_ LiMP types, and an open-ended number of *n* LiTT types (one for the life time of each movement group type to which an individual can belong). Horizontal whiskers on icons represent variations in the length of examples of the same type within a category (except for metaFuMEs which are the length of time between successive relocation points and DARs which are fixed to the biological relevant diel cycle) and vertical whiskers indicate some building-block variation within the same type (except for FuMEs which are much more stereotyped, and hence much less variable within types than other segmentation categories). Color shades within categories represent different types within those categories. Colorless strings of icons indicate how each category can be considered as a string of the next lower category elements (i.e., shapes and general color denote hierarchical level, while different shades of the same color indicate within level variation). Because FuMEs are hard to identify using relocation data only, a hierarchical segmentation of lifetime tracks will typically be supported by a metaFuME baseset rather than by a set of FuMEs themselves.

Segmenting relocation data at a metaFuME level only makes sense if the data have been sampled at frequencies that lie somewhere between the time it takes to execute several typical FuMEs (i.e., on the order ten seconds) and the time it takes to perform a short-period CAM that is relevant to movement ecologists (i.e., on the order of a few minutes, except for sharp burst of movement related to predation or defense). Fortunately, current technologies facilitate the collection of data at frequencies of 0.01 to 1 Hz in birds (Harel, Horvitz & Nathan 2016, Harel & Nathan 2018) and mammals (de Weerd et al. 2015, McGavin et al. 2018) of even relatively small size. Beyond GPS methods for collecting high frequency data are other methods such as reverse GPS (e.g., the Atlas system; Weiser et al. 2016, Toledo et al. 2016); but reverse GPS is generally limited in spatial extent to some tens or hundreds of square kilometers. Also, high-frequency movement data has been collected using video equipment for small (e.g., ants) to moderately-sized (e.g., mice) organisms in a laboratory setting (Spink et al. 2001, Kane et al. 2004, Delcourt et al. 2013). From a practical point of view, though, particularly since the size of relocation data sets can be rather large when collected at frequencies of around 0.01 to 1 Hz, it may be useful to carry out a metaFuME identification process on selected segments of the full path *W* (such as CAMs), and then use a suitable method to reconcile disparate identification efforts to obtain a consensus set of metaFuME types that can be applied to the full movement path.

Assuming that a set of metaFuMEs has been identified, as discussed in more detail in succeeding sections, then it may be possible to stably identify different kinds of short duration CAMs with periods varying from around 10 metaFuMEs (i.e., a couple of minutes) to a few hours (Fig. 1, Table 1). Following this, longer duration CAMs may be more conveniently parsed in terms of several shorter duration CAMs rather than in terms of metaFuME types. To emphasize this, both short and long duration CAMs are identified in the hierarchical scheme laid out in Table 1. Due to the importance of circadian rhythms as physiological and behavioral drivers (Takahashi et al. 2008, Yerushalmi & Green 2009, Hardin & Panda 2013), the diel cycle provides an empirically obvious segmentation window for the identification of various types of diel activity routines (DARs; set of green boxes in Fig. 1; also see Table 1). Several types of DARs may then be identified in terms of differences in the type and frequency of their constituent CAMs. For example, different types of DARs have been identified in terms of the portion of their diel cycle that various groups of lemurs are active (Donati et al. 2016) and of the variation in the diel travel rates of turtles (Jonsen et al. 2006).

DARs, in turn, can be strung together to create life-history movement phases (LiMPs; sets of red pentagons in Fig. 1) that, depending on the longevity of the species, may be periodic (e.g., annual migration; for a review see migration Milner-Gulland et al. 2011) or episodic (e.g., dispersal; for a review see Bowler & Benton 2005). Finally, a sequences of LiMPs sequentially strung together from the birth-to-death of an individual constitute its full life time track (LiTT; Nathan et al. 2008) (indicated by the grey pentagons in Fig. 1; also see Table 1). Beyond LiMPs, LiTTs from several individuals can be used to map out space-use by populations rather than just the movement path of any single individual (Mueller & Fagan 2008).

This pragmatic view of metaFuME, CAM, DAR and LiMP segmentation of lifetime tracks, as illustrated in Fig. 1, provides a hierarchical classification system of appropriate complexity (Getz et al. 2018, Larsen et al. 2016) for the parsing of movement paths based on two different fixed time-segment scales: metaFuMES (sampling interval at the order of HZ or 0.1 HZ) and DARs (24 h). The metaFuMEs provide a basis for constructing CAMs as the biologically meaningful segments of DARs; and DARs are used to support the construction of seasonally or life-history relevant LiMPs. Given this ecological relevance of canonical activity modes and lifetime tracks, the classification system presented here also provides a semantic framework for the generation of quantitative narratives (Getz, Salter & Tallam 2019) needed to explain the role that movement plays in the ecology and evolution of organisms. In the context of evolution, for example, we expect natural selection to hone both the mechanics of performing FuMEs and the way these FuMEs are organized into metaFuMEs, CAMs, DARs, and LiMPs to create lifetime tracks that maximize the expected lifetime fitness of individuals. In particular, it is of considerable interest to assess how the frequency and sequencing of different FuMEs and CAMs in the production of DARs and LiMPs affect individual fitness (Hein et al. 2016, Cattarino et al. 2016).

Beyond the relevance of diel cycles to movement behavior (Wittemyer et al. 2008), are lunar (Polansky et al. 2010) and seasonal cycles (Marra et al. 2015); and even weekly cycles if the influence of humans has some impact on the movement of animals in urban and suburban areas. The critical nature of the diel scale in driving movement behavior is reflected in our existence of the fixed-period DAR category in framework presented in Fig. 1. In contrast, the lunar cycle may be more relevant to some organisms than others (Polansky et al. 2010). Thus beyond DARs only seasonally or life-history relevant LiMPs segments are identified with seasonal segments only being relevant to organisms with life-spans long enough to experience several seasonal cycles (Marra et al. 2015, Allen et al. 2018). For some organisms, particular LiMPs, such as dispersal, occur only once; while migration behavior may vary from year to year, influenced by inter-annual variations in climatic conditions—perhaps linked to multiyear marine (Mysterud et al. 2001, Grémillet & Boulinier 2009) or sunspot (Myers 2018) cycles. Further, periodically driven movement patterns may also change with age (e.g., exploration when young) and life-history stage (e.g., elephants in musth). Thus, beyond DARs, other cyclic patterns become either less obvious or more species-specific.

## The Data Compatibility Problem

Parsing a movement path into its elements is somewhat like picking out words from a voice recording. The human brain does this very well, as do modern digital machines using deep-learning methods (Hinton et al. 2012, Zhang et al. 2018). In an extremely crude sense, FuMEs are comparable to syllables, metaFuMEs to words, short duration CAMs to sentences and long duration CAMs to paragraphs. DARs may be thought of as single pages in a book-of-life, where each page can be identified as belonging to one of a rather limited number of types (e.g., a typical winter versus summer day or a day during a migratory versus non-migratory LiMP). Thus, the stories being told are rather boring, page-by-page repetitions, with important cyclic variations, as well as other variations due to environmental influences.

Despite the current, technology driven, exponential growth in the quantity and quality of movement data (including relocation, accelerometer, physiological, and environmental measurements; Williams et al. 2019), collecting the kind of multi-body-placement accelerometer and relocation data needed to identify the start and finish of individual FuMEs (i.e., with a spatial accuracy of centimeters rather than meters or even decimeters) may still be beyond the budgets of most movement ecology studies. Other technologies, such as audio (Sapir et al. 2010, Hurme et al. 2019, Northrup et al. 2019), magnetometer (Williams et al. 2017, Chakravarty et al. 2019), and video (Spink et al. 2001, Kane et al. 2004, Delcourt et al. 2013) may well be more cost effect in terms identifying the start and finish times of individual FuMES. Thus, for some time to come, relocation data collected at fixed frequencies slower than around 1 Hz are going to be significantly misaligned in time (at least to within half a second) with the start and finish of each of the FuMEs that make up a path segment of interest.

At this point, it is worth noting that accelerometer data alone has been used to parse out relatively short time scale (order of 10 secs) behavioral elements along movement paths using machine learning methods (Nathan et al. 2012, Thessen 2016, Wang et al. 2015). These behavioral elements, which almost certainly include several FuME steps, in reality are partial elements of more extensive short-duration CAMs that typically last tens of seconds to several minutes. Accelerometer data, for example, has been used to categorize standing, running, preening, eating, and active and passive flight CAMs in vultures with 80-90% accuracy (Nathan et al. 2012). Another analysis on baboons succeeded in using accelerometer data to categorize foraging, lying, sitting, running, standing, walking, groom (actor), groom (receiver) with high accuracy (*>* 90%), except for grooming, which was considerably harder to identify, particularly when the focal individual was the receiver (Fehlmann et al. 2017). In pumas, accelerometer has been used to identify low versus high acceleration movement phases, resting, eating and grooming (Wang et al. 2015). In these examples, sitting/resting, grooming, preening and feeding CAMs are identifiable, but none of the basic FuMEs executed during the generation of these CAMs are individually identified. It may be possible for some FuMEs, however, such as the time taken to complete a FuME when walking versus running, to be identifiable directly from accelerometer data (as we find in modern digital watches that are able to monitor the number of steps we take while moving).

In summary, location data on its own (i.e., without accompanying subsecond accelerometer, acoustic, magnetometer or video data) is fundamentally incompatible with the identification of FuMEs because FuMEs are characterized by the movement of body parts while location data apply to a body as a whole. For this reason, it is worth reemphasizing the following.

1. Given that the identification of individual FuMEs from relocation data is not generally possible, we are left with the rather challenging task of extracting a set of metaFuME elements, where each metaFuME type is characterized by its own SL-TA distributional pair and correlations with and between consecutive and simultaneous values.
2. The complete set of metaFuME types, once identified, can then be used as a basis for constructing an appropriate complexity (Getz et al. 2018, Larsen et al. 2016), hierarchical frame-work of CAM, DAR and LiMP segments, where the latter is used to classify different types of lifetime tracks (Table 1), beginning with CAM sequences constructed using a multi-CAM metaFuME Markov (M^3^) modeling approach described in more detail below.

## Current Path Analysis Methods

Before considering how to find a set of metaFuME distribution set that provides an appropriately defined “best fit” to an empirical walk *W* represented in 2-D, by the relocation time-series

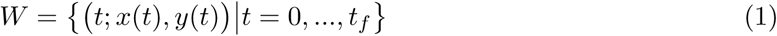

let us briefly review the current state of path analysis that is relevant for analysing relocation data sampled at scales typically appropriate for identifying CAMs and higher-level patterns (Fig. 2). Note that the temporal resolution of the data will determine whether it is possible to identify short duration CAMs (e.g., vigilance behavior during grazing; Fortin et al. 2004), long duration CAMs (e.g., heading to water; Polansky et al. 2015), or only movement patterns at diel scales and beyond (Spiegel & O’Farrell 2019, Owen-Smith 2013, Giotto et al. 2015).

At any time scale, movement time series may be analysed using purely statistical methods applied to the SL and TA time series derived from *W* to generate various running statistics. These statistics may include the means, variance, autocorrelations of SL and TA time series data and cross correlations between the two (Chatfield 2016, McCulloch & Cain 1989, Bergman et al. 2000, Byers 2001). Running versions of these statistics can then be used to identify points in time where abrupt changes in their values occur, using methods referred to as behavioral change-point analyses (BCPAs; Owen-Smith & Martin 2015, Chen & Gupta 2011, Matteson & James 2014, Killick & Eckley 2014, Gurarie et al. 2009, 2016; Fig. 2) or, more generally, path segmentation methods (PS; Nams 2014, Edelhoff et al. 2016, Seidel et al. 2018).

**Figure 2:**
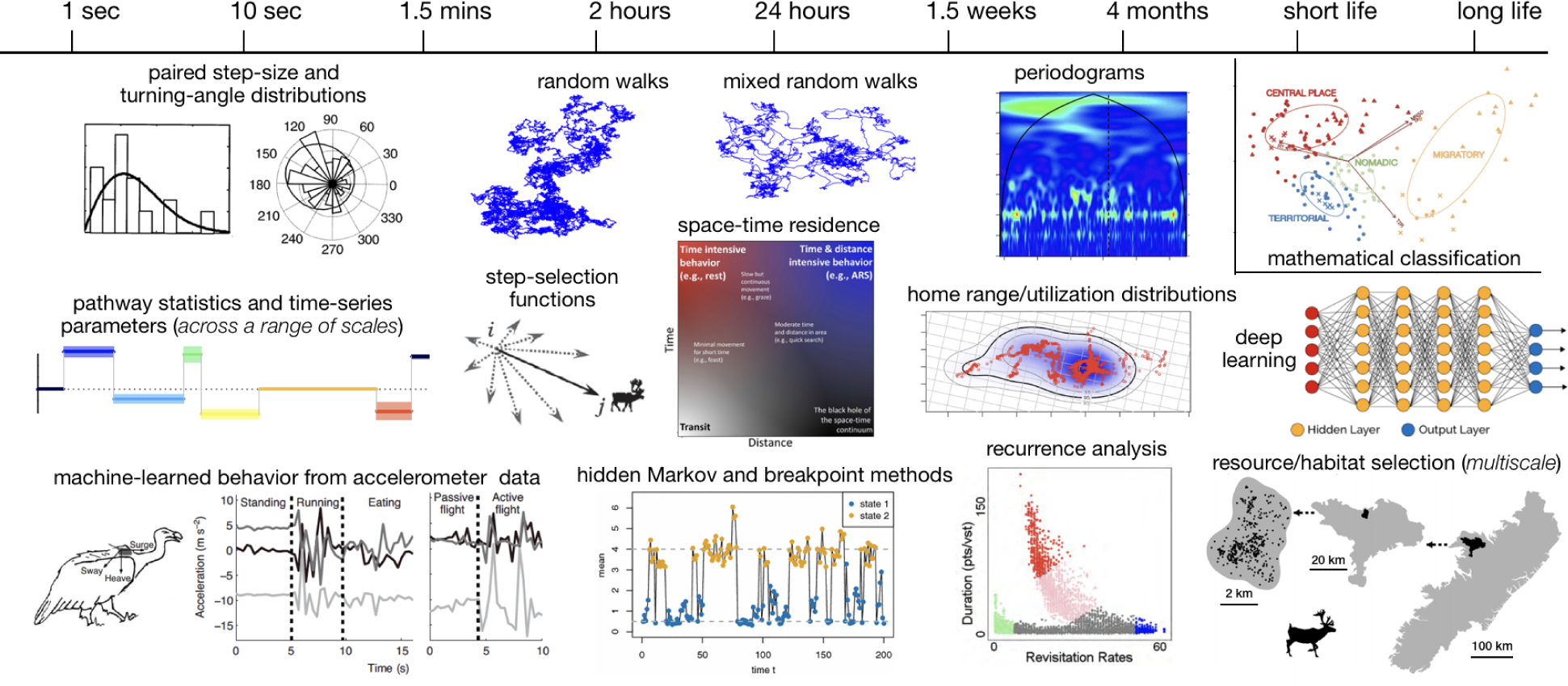
Current scale-dependent analytical methods for analysing movement paths. The temporal range is only suggestive and best applied to medium and large vertebrates. Also the image placements are not precise and some methods, such as machine learning (Thessen 2016), can be applied to data at any scale, but here are associated with the scale at which they are likely to be most useful. In addition, deep learning (useful for identifying different types of long-term patterns) is actually a subset of machine learning (where other deep-learning techniques, such as random forests and support vector machines have been applied to accelerometer data; Nathan et al. 2012, Fehlmann et al. 2017). Stochastic walks include correlated and biased random walks (Johnson et al. 2008). Space-time residence analyses represents a family methods that include first-passage-time (FTP; Fauchald & Tveraa 2003, McKenzie et al. 2009) and related approaches (Torres et al. 2017), while recurrence analyses cover a plethora of methods used to identify recursive movement patterns (Berger-Tal & Bar-David 2015, Bar-David et al. 2009). The melange of images is extracted from publications in the literature (Nathan et al. 2012, Panzacchi et al. 2016, Morales et al. 2004, DeCesare et al. 2012, Wittemyer et al. 2008, Lyons et al. 2013, Abrahms et al. 2017, Fleming & Calabrese 2017, Pohle et al. 2017, Torres et al. 2017, Gurarie et al. 2017), as well as created for this publication.

Beyond purely statistical characterizations of the SL and TA time series is the notion that some sort of state (typically behavioral; but possibly physiological, disease, etc.) can be associated with each location point in the walk *W* (Patterson et al. 2008). A very simple example is the stop/move representation of cattle grazing paths (Zhao & Jurdak 2016). The most widely used methods for identifying subsets of points that reflect some underlying state or canonical activity mode (CAM) involve hidden Markov models (HMM) (Michelot et al. 2016, Franke et al. 2004, Langrock et al. 2012, Zucchini et al. 2016) (Fig. 2), although an independent mixture modeling approach has been shown to be equally viable, if not better (Goodall et al. 2017). These methods are particularly germane when covariate data are available at each location, such as accelerometer and temperature data from GPS collars, remotely sensed physiological information (Cross et al. 2016), vegetation structure (Tsalyuk et al. 2019), or physical landscape features that, for example, facilitate or retard movement (Zeller et al. 2012). The task of an HMM is to find a best estimate of which state of a current set of “hidden” states *H*_*i*_ belonging to a set of *n* possible states 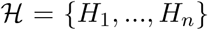 should be assigned to the individual when at the location (*x*(*t*), *y*(*t*)).

HMMs can be applied at any temporal scale, including the metaFuME scale, depending only on the relocation and covariate data. Other methods, such as step-selection function (SSF) methods, which address the question of where am I likely to move next as a function of landscape covariates, can be applied at any scale (Fortin et al. 2005, Panzacchi et al. 2016, Thurfjell et al. 2014, Avgar et al. 2016). SSFs, however, address different kinds of questions at different scales: they have more of a behavioral interpretation at a 1-minute scale while more of an ecological (e.g., resource use) interpretation at multi-hour scales. On the other hand, habitat or resource-selection function (HSF or RSF) methods (Boyce et al. 2002, McGarigal et al. 2016, DeCesare et al. 2012) are, by their very nature, low resolution models that typically apply at a seasonal level and assume non-autocorrelated location data. When RSFs are evaluated at a lifetime level over many individuals in a population, they are called niche models—but such models no longer rely on movement data, only on point location data (Townsend Peterson et al. 2007).

Other general time-series or signal-analysis methods that have been co-opted by movement ecologists include wavelet analysis (Wittemyer et al. 2008, Polansky et al. 2010)—which provides interpretations at the resolution of one to several orders of magnitude greater than the frequency at which data are collected (e.g., daily movement to monthly movement patterns for data collected at the frequency of 1-hour for at least a year)—and machine learning approaches (Thessen 2016, Valletta et al. 2017) that have been applied both to location (Bennison et al. 2018) and accelerometer data (Nathan et al. 2012) (Fig. 2). These methods, however, do not have an extensive history of application in movement ecology, but are likely to become more ubiquitous as the field of movement ecology matures.

## Stochastic Walk Statistics

As discussed earlier, identifying a set of FuMEs is a problem that requires biomechanical (Delp & Loan 2000), audio (Sapir et al. 2010), or other kinds of covariate data (Chakravarty et al. 2019) than relocation data on its own. With relocation data alone, a hierarchical analysis of a path can only be underpinned by elements that are derived statistically from the relocation time series *W* (Eq.1).

From this time series we can extract a 1-D time series of step lengths and another 1-D time series of turning angles as follows:

- Generate the step-length (SL) time series

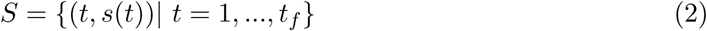

using the derived values

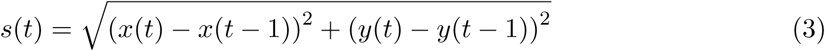
- Generate the turning-angle (TA) time series

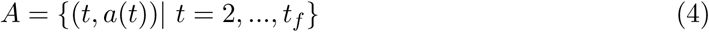

using the derived values

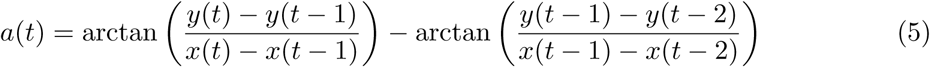

From these two time series we can obtain their means and variances respectively denoted by (*µ*_*ℓ*_, *σ*_*ℓ*_), *ℓ* = *s, a*, and then use them compute the following three variation related time series:

- Generate two normalized (by the appropriate variances) “running-term autocorrelation” time series

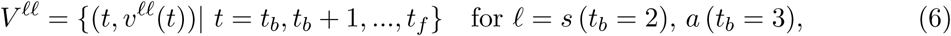

where

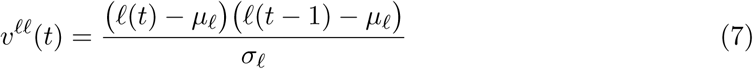
- Generate a normalized “running-term cross-correlation” time series

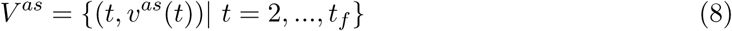

where

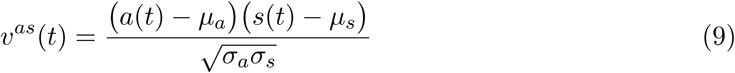

We are now challenged with the task of using the five time series above (step length, turning angle, two auto correlation and one cross correlation)—and, perhaps, other covariate data when available (e.g., acoustic, accelerometer, local environmental)—to parse *W* (Eq. 1) into several, say Λ, different single-step metaFuME ensembles (Fig. 3). The most obvious approach is to use clustering methods, but this should be accompanied by some sort of range standardization procedure for each variable to improve the performance of the clustering algorithm (Van Moorter et al. 2010). Each of these Λ ensembles, belonging to the same metaFuME type, can now be regarded as a set of drawings from the same bivariate distribution *D*_*λ*_(*s, a*), *λ* = 1, …, Λ. If the relocation sampling period, except for speed bursts, encompasses around 10-50 FuMEs (i.e., more than around 5 seconds but less than about a minute—e.g, fast walking in humans crudely corresponds to 2 steps per second, though most movement is slower than this); then, in the proposed hierarchical segmentation framework, we are in the metaFuME segment zone (Fig. 1, Table 1). If the relocation sampling period is on the order of tens of minutes to several hours, then each relocation likely encompasses hundreds to tens of thousands of FuMEs. In this case, set of states are more appropriately identified as short or long duration CAMs than as metaFuMEs.

**Figure 3:**
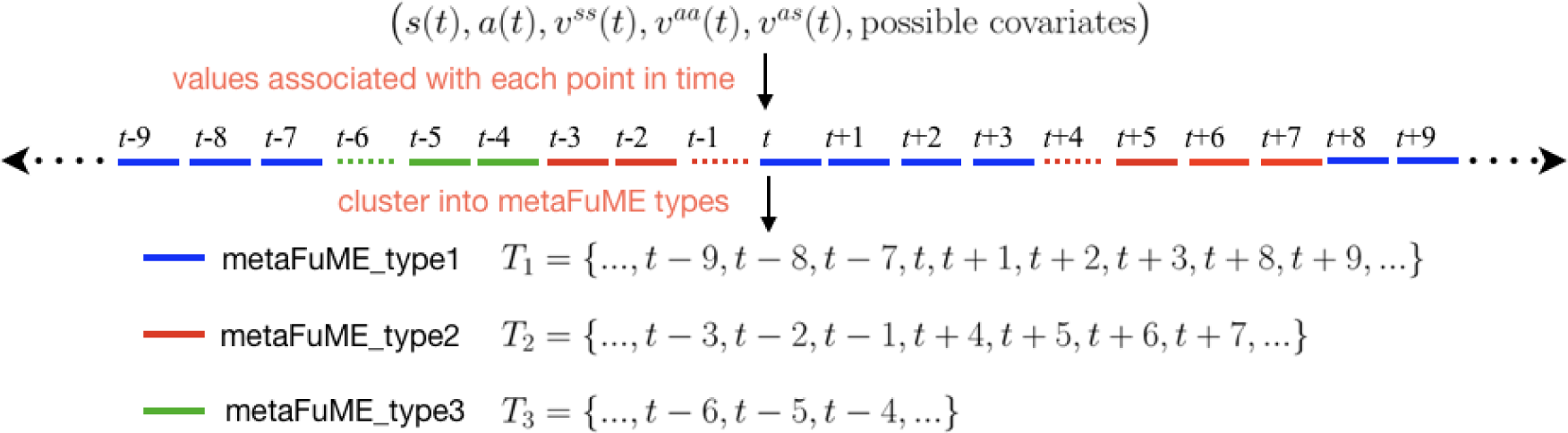
Parsing the walk *W* into a set of metaFuMEs. Associated with each point *t* is a set of running values (*s*(*t*), *a*(*t*), *v*^*ss*^(*t*), *v*^*aa*^(*t*), *v*^*as*^(*t*), other possible covariates). Cluster analysis may be used to generate metaFuME categories. For purposes of illustration, we depict three types of metaFuMEs with associated lists of time steps—i.e., metaFuME sequence index sets—that can then be used to define the ensemble of values associated with each metaFuME type: blue = *T*_1_, red = *T*_2_, and green = *T*_3_. Note that we expect transition points between strings of metaFuMEs of one type (broken line segments), such as *t* − 6, *t* − 1 and *t* + 4, could well be outliers in the clustering process. Outliers may be assigned to the cluster to which they are closest in some suitably defined sense (as indicated by the colors of the broken lines).

Once a set of metaFuMEs has been identified from a particular ensemble of segments of the same CAM or DAR type, along with the accompanying metaFuME sequence index sets (Fig. 3), we are still faced with the task of reconciling various metaFuME sets obtained from different CAM or DAR ensembles into a comprehensive metaFuME set. Such a set will then underlie all LiMPs of a particular type, but not necessarily across a complete lifetime track, because we might expect metaFuME sets for, say, juveniles versus adults to be different for same species. Many different types of cluster analyses (McGarigal et al. 2016) can be tried to identify metaFuMEs. Also, multicriteria optimization methods using, for example, genetic algorithms or approximate Bayesian computations (Coello 2003, Odu & Charles-Owaba 2013, Marin et al. 2012) could be used to reconcile metaFuME sets obtained from ensembles of different types of segments. Such investigations, however, are well beyond the scope of this “concepts” paper, but are undertaken elsewhere (Getz, Vissat & Salter 2019).

Once the time series *W* and, hence, the corresponding SL and TA time series have been parsed into ensembles and organized into a set of Λ metaFuMEs, the associated metaFuME sequence index sets *T*_*λ*_, *λ* = 1, …, Λ, (Fig. 3) can be identified. The distributions of all the joint pairs (*s*(*t*), *a*(*t*)), where *t* ∈ **T**_*λ*_, *λ* = 1, …, Λ, can then be fitted to an appropriate metaFuME bivariate distributions *D*_*λ*_(*s, a*), *λ* = 1, …, Λ and the metaFuME sequence index set ensemble 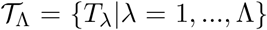 can be used to estimate transition rates among metaFuME sequences. In essence, we have estimated a set of bivariate metaFuME distributions

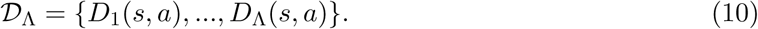

and a matrix of values (**P**)_*λκ*_ = *p*_*λκ*_ (with *λ*, *κ* = 1, …, Λ) that represent the probability of sampling values from the distribution *D*_*λ*_(*s, a*) when the previous drawing was from the distribution *D*_*κ*_(*s, a*).

We can now use the set of distributions *𝒟*_Λ_ and transition matrix **P**, essentially as a Markov metaFuME movement model, to construct a CAM segment of a walk *W* (*t*_*f*_). We do this by first generating a set of *T* drawings (the hat notation represents drawings)

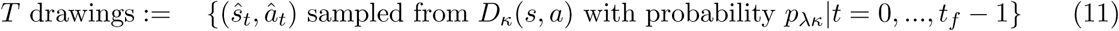

and then generating a segment of *W* (*t*_*f*_) from these drawings using the following equations (Patterson et al. 2008), where *θ*(*t*) is the absolute angle of heading at time *t*

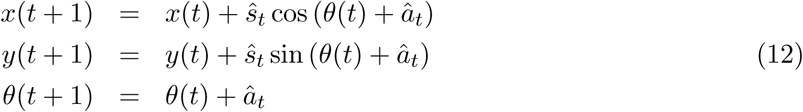

Thus, in short, we use the extracted identified metaFuME-distribution-set and Markov transition-matrix pair (*𝒟*_Λ_, **P**) to generate a CAM segment of a walk *W* that is one instantiation of an ensemble of CAM segments generated using Eqs. 11 and 12.

To generate different sets of CAM sequences and string these together to form particular DARs requires that we have additional distributional descriptions for the length of different CAM sequences found in particular DARs, and matrix probabilities for transitioning among CAMs within such DARs. The outlines of model to accomplish this task, referred to as a multi-CAM metaFuME Markov (M^3^) model is described in the next section. In terms of a single CAM metaFuME Markov model embodied in Eqs. 11 and 12, the appropriate number of metaFuME distribution pairs Λ in the set *𝒟*_Λ_ (Eq. 10) is, *a priori*, unknown. A first guess at this number might be the number of modes in step length time series derived for different types of CAM segments, if such modes are evident. The problem of finding the best mixture of unimodal pairs 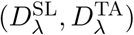 of distributions that best fit such derived empirical data is a challenging estimation problem that can be approached in several ways, including ensemble Kalman filtering (Dovera & Della Rossa 2011) under the assumption that the underlying distributions are log-concave (Walther et al. 2009). A prudent approach may be to proceed by first looking for the two best-fitting pairs of bivariate distributions:

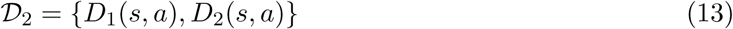

and then moving on to three, four, and so on, until no improvement in fit is obtained at the next value for Λ in an information theoretic sense (Symonds & Moussalli 2011). We may then use existing segmentation methods (e.g., BCPA) to identify possible CAMs that may emerge from the simulated data and compare them to the CAMS identified in the original data If the fit is satisfactory, we may then use our multi-CAM metaFuME Markov (M^3^) model (Getz, Vissat & Salter 2019) to predict how individuals may respond to management or global change in terms of local and intermediate scale movements. We note, though, that the M^3^ modeling approach described in the next section contains no global directional information. Thus, additional constructs will be needed to obtain movement motivated by headings to specific distant target sites. The performance of M^3^-type models and their extensions are investigated elsewhere (Getz, Vissat & Salter 2019).

## Construction of M^3^ Model

The basis of the M^3^ model is to take ensembles of CAM segments of the same type and identify a set of metaFuMEs that can be used to simulate the local structure of such CAMs. If this is done for all the kinds of CAMs that constitute a particular DAR type, then the model can be used to string several types of CAMs together to conform to observed sequences and frequencies of CAMs characteristic of type of DAR under consideration. This approach to constructing M^3^ models has been explored and evaluated elsewhere (Getz, Vissat & Salter 2019). Such M^3^ models, however, do not account for movement towards specific distant targets. To include such phenomena requires that M^3^ models be appropriately extended to include directional biases induced by an attraction to distant geographic locations or a repulsion related to the existence of boundaries to movement (e.g., landscape topography or water).

Extraction of an M^3^ model from relocation data requires that the frequency of the data be available at the metaFuME scale, which from Table 1 for medium to large terrestrial vertebrates is around 0.01 to 0.1 Hz. Although details of the approach can be found in (Getz, Vissat & Salter 2019), for the sake of completeness, a brief summary of the approach is enumerated here.

### Summary of approach

1. Path relocation data, *W*^data^ (Eq. 1) is parsed into ensembles of DARs of various types, using an appropriate method (e.g., based on net-square daily displacement or other suitable daily measures; see Bunnefeld et al. 2011, Bischof et al. 2012, Abrahms et al. 2017, Owen-Smith & Martin 2015, Owen-Smith et al. 2010).
2. CAM segments are identified from DAR segment ensembles of the same type (Fig. 4), using appropriate methods such as BCPA (Nams 2014, Edelhoff et al. 2016, Seidel et al. 2018) or HMM (Zucchini et al. 2016). In the latter case, the HMM will need to be performed on a subset of the data subsampled at a scale suitable for identifying CAMs of interest (e.g., if CAMs of interest are assumed to last around 15 minutes or longer then data should be subsampled using a consecutive point intervals of, say, around 5 to 10 minutes).
3. All the data from the ensemble of path segments identified as belonging to the same CAM type within the same DAR type should be strung together into a set of points. If we assume that Φ such CAM-DAR compound types are identified, use 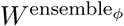, *φ* = 1, …, Φ to denote these sets of points.
4. Cluster analyses or other type of categorization procedure should be performed, using the values (*s*(*t*), *a*(*t*), *v*^*ss*^, *v*^*aa*^, *v*^*as*^) (Eqs. 2–9) for all times *t* for which there are points in the sets 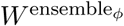 to identify sets of metaFuME types specific to each of the sets 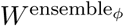, *φ* = 1, …, Φ (Fig. 3).
5. For each ensemble type *φ*, distributions 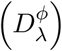, *λ*_*φ*_ = 1, …, Λ_*φ*_, should be fitted to each of the Λ_*φ*_ metaFuME types identified in the previous step, with some effort to reconcile metaFuME types identified across different ensembles 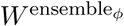, *φ* = 1, …, Φ
6. Distributions 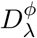, *λ* = 1, …, Λ_*φ*_ can be fitted to each of the metaFuME types identified in ensemble *φ*, *φ* = 1, …, Φ, and an associated Markov transmission matrix (**P**_*φ*_) estimated to obtain the characterizing pairs (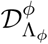, **P**^*φ*^), which can then be used in a Markov metaFuME movement model to generate CAMs of type *φ*, for *φ* = 1, …, Φ.
7. The metaFuME Markov models derived in the previous step for each type of CAM can be combined into a multi-CAM metaFuME Markov (M^3^) model that produces sequences of CAMs with CAM lengths and CAM-type transition statistics that are extracted from the DAR ensemble data.
8. The performance of the M^3^ model can be evaluated by comparing the CAMs obtained directly from *W*^data^, using existing BCPA and HMM methods, with CAMs produced by the M^3^ model simulations.

**Figure 4:**
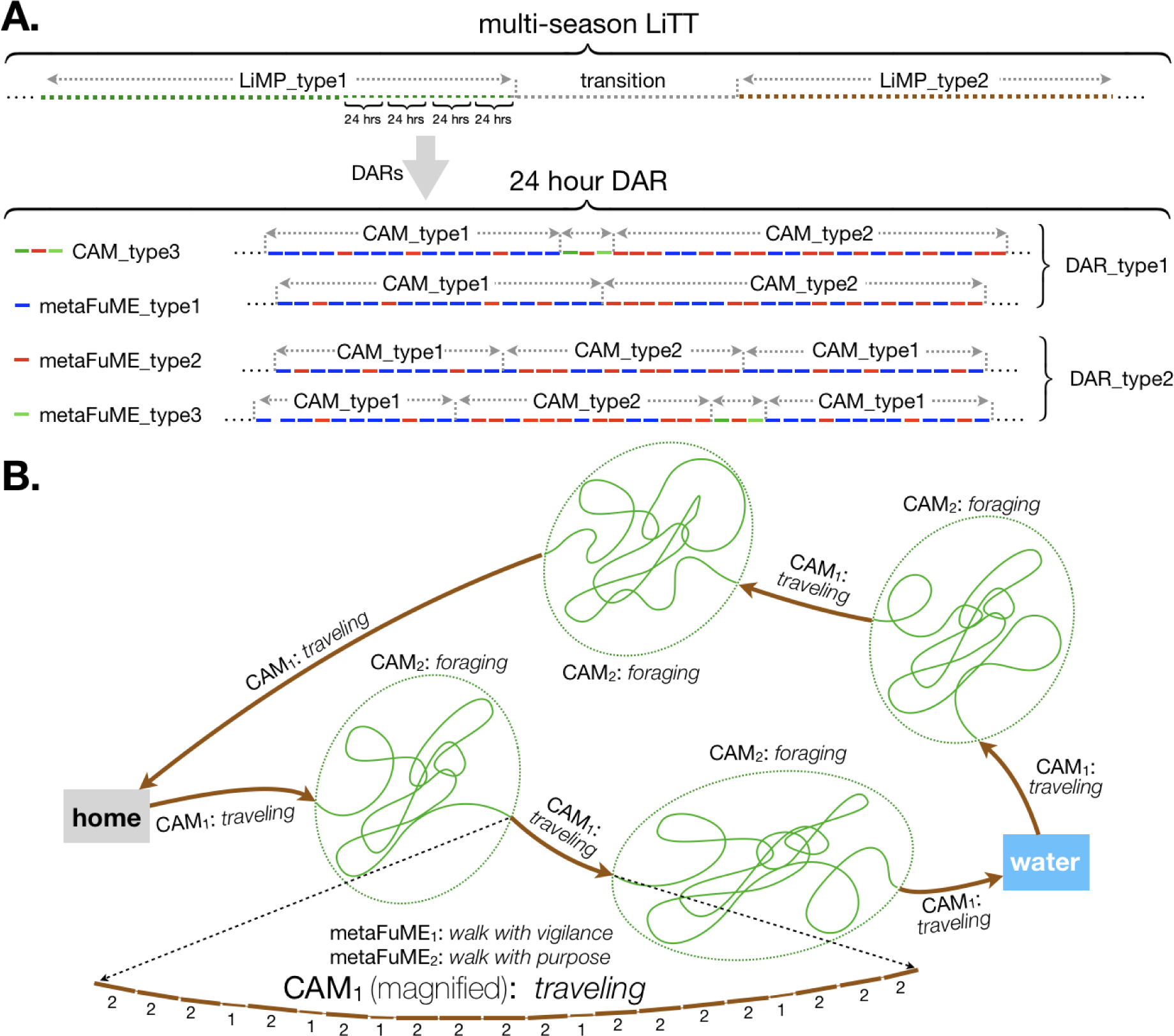
The segmentation structure of movement paths at different scales. **A.** A multi-season life time track (LiTT) *W* (Eq. 1) can be segmented into different types of life-history movement phases (LiMPs), each of which can then be segmented into diel activity routines (DARs) and again segmented into canonical activity modes (CAMs). Each CAM is constructed from different proportions of metaFuMEs (aggregates of correlated fundamental movement elements), where each metaFuME is derived from a bivariate step-length/turning-angle distribution derived from time series with specific first-order correlative statistics. Short-duration CAMs (e.g., CAM 3) can easily be missed and averaged into longer duration CAMs (e.g., CAMs 1 and 2), if DAR segmentation analyses is not undertaken at a sufficiently fine scale. **B.** An illustrative example of a simple two-CAM *return-to-home-with-a-water-stop* DAR where the two types of metaFuMEs that generate the *traveling* CAM are shown under magnification.

## Conclusion

Without an underlying framework to organize information, deep understanding within any scientific field is impossible. Clearly this has been true of the physical sciences with theories of gravity and the structure of matter being central to our current understanding of the physical universe. This is also true of the life sciences where, for example, the Linnaean taxonomic system, with some modifications, has reigned supreme over the past 250 years (Schuh 2003). This reign still holds today, even within the microbial sciences (Rosselló-Móra & Whitman 2019, Moore et al. 2010) where Mayr’s biological species concept fails in several critical ways (Gevers et al. 2005).

In the context of movement ecology, the classification of movement types at different spatiotemporal scales is of considerable interest, although approaches to date have been somewhat informal. At the lifetime track level, basic life-history types regarding, *interalia*, dispersal and migration behavior may be identified. In the movement ecology literature, we see many studies interested in diel activity routine (DARs) (Rahimi & Owen-Smith 2007, Owen-Smith 2013, Owen-Smith & Goodall 2014), lifetime phase (LiMPs) (Fahr et al. 2015, Marra et al. 2015) and over-all life time tracks (LiTTs) (Abrahms et al. 2017). Contrasting DAR types may include distinctions among nomadic, central-place foraging, or territorial behavior, where a single lifetime track may have movement phases dominated by one or other daily activity routine, as in male springbok in Etosha National Park, Namibia, exhibiting territorial behavior during the wet season and nomadic behavior during the dry season, with daily excursions to the same water-hole around midday during the wet season (Lyons et al. 2013). Thus daily activity routines are also very likely to be influenced by seasonal factors, as in pandas that change diel levels of activity in response to seasonal drivers (Zhang et al. 2017).

Extensive effort has also been made to parse diel and longer segments into various types of 4 sub-diel activity modes (Owen-Smith et al. 2010, Donati et al. 2016) that, if stably identifiable across different segments, can be organized into a set of canonical activity modes (CAMs; Getz & 4 Saltz 2008). At finer time scales, however, beyond using accelerometer, acoustic and magnetometer 4 data to identify behavioral states (Williams et al. 2017, Hurme et al. 2019, Nathan et al. 2012, Fehlmann et al. 2017, Chakravarty et al. 2019, Sapir et al. 2010, Zucchini et al. 2016), very little work has been undertaken to identify sets of fundamental movement elements (FuMEs; same as FMEs in Getz & Saltz 2008) from which CAMs are constructed. As pointed out in this paper, using relocation data alone, we cannot endeavour to identify an underlying set of FuMEs. The best we can hope for, provided subminute relocation data is available, is the identification of a set of metaFuMEs consisting of repeated or correlated strings of FuMES, and characterized by particular step-length and turning-angle distributions.

We should not underestimate the analytical and computational challenge required to extract a comprehensive and stable (across many segments at various scales) set of metaFuME elements. The best methods to do this still remain to be developed and how well this may be accomplished remains to be seen; although satisfying progress has already been made (Getz, Vissat & Salter 2019). Future progress may require many of the ideas presented here to be refined or modified. There is not denying, however, that formalized, widely accepted, hierarchical-path-segmentation framework will greatly facilitate efforts to address outstanding questions in movement ecology, particularly those involving comparative analyses within species (Wittemyer et al. 2019), as well as across species. Within species variation may allow us to assess differences along geographic clines with application to the behavioral adaptation of species under landscape and climate change (Seebacher & Post 2015). It may help us assess fitness in the context of feeding strategies, social behavior (Harel et al. 2017, Harel, Duriez, Spiegel, Fluhr, Horvitz, Getz, Bouten, Sarrazin, Hatzofe & Nathan 2016), areas attractive to populations (Giotto et al. 2015), or mapping out landscapes in terms of their overall resistance to movement (Zeller et al. 2012). It may also be diagnostic of changes in movement behavior when individuals are stressed, ill, have genetic defects, or females are pregnant (Spink et al. 2001, Owen-Smith 2013); or it may be predictive in terms of pathogen transmission and the spread of disease (Cross et al. 2007, Tracey et al. 2014, Dougherty et al. 2018, Zidon et al. 2017). Differences in various species’ movement profiles at various scales may be critical when it comes to taking a multispecies approach to assessing the impacts of ecosystem management on species conservation (Brodie et al. 2015, Runge et al. 2014).

Additionally, a formalized, hierarchical-path-segmentation framework of appropriate complexity (Getz et al. 2018, Larsen et al. 2016) can be used, for example, to address the dozen plus questions that were recently posed regarding the movement ecology of marine megafauna (Hays et al. 2016)— but are equally applicable to all animal species. These questions include: 1) “Are there simple rules underlying seemingly complex movement patterns and hence common drivers for movement across species?” and 2.) “How will climate change impact animal movements?” An ability to address both of these question, as well as those raised above and others besides, in a comparative way—with a level of consistency and coherency that only a universally accepted classification framework can provide—is needed with great urgency, as the field of movement ecology matures. The framework presented here is a starting point to developing a more comprehensive one that can be fleshed out in the future, supported by analytical and computational methods, some of which still need to be developed or refined from existing methods. The framework proposed here, however, provides a guide to constructing M^3^ models for simulating CAM and DAR segments in empirical relocation data (Vissat et al. 2019), with extensions yet to be developed, will ultimately provide a model for simulating walks that conform to empirical data at several local (metaFuME, CAM) and global scales (biased CAM components in DARs and beyond). It will take models of this complexity to address the two questions posed here, as well as those posed elsewhere in the context of theoretical and applied movement ecology, the latter in particular addressing questions related to conservation biology, resource management, and global change assessment.

## Glossary

An expansion of the various acronyms used in the text are provided here in alphabetical order along with a very brief definition/explanation of their meaning.

**BCPA:** Behavioral Change Point Analysis. Behavioral change-point analysis refers to a group of methods used to determine how the statistics of a biological variable *y* (e.g., its mean, variance, autocorrelation, or rate of change in slope or curvature), dependent on a second variable *x*, switches at threshold points with changes in *x* (e.g., space, time, or the abscissa in a stress-response relationship) (Morales et al. 2004, Andersen et al. 2009, Chen & Gupta 2011, Jonsen et al. 2007). Note, more generally, *x* could be vector-valued.

**CAM:** Canonical Activity Mode. This is a stably classifiable subdiel behavioral mode (pattern of movement) such as a foraging bout, resting period, or purposeful heading (i.e., traveling) to a distant target location.

**DAR:** Diel Activity Routine. This is a stably classifiable 24-hour sequence of CAMs that occur at characteristic frequencies and times of the day.

**FuME:** Fundamental Movement Elements (also FME; Getz & Saltz 2008). This is a relatively rapid, highly repeatable, stereotypic set of body movements that forms the basis of the locomotory capacity of an individual (e.g., a walk step, a running step, a wing flap, a jump, etc.)

**HMM:** Hidden Markov Model. A time series of observable values that depend in a probabilistic sense on the values of an associated, but unobservable Markov chain process (a set of states where the transition from one state to another depends only on the value of the current state) (Zucchini et al. 2016)

**LiMP:** Life-history Movement Phase. This is a path segment that typically reflects a life-history relevant movement behavior such as dispersal (episodic), migration (periodic), or other periodic behaviors at a greater-than-diel scale.

**LiTT:** Life Time Track. This is the total movement path of an individual from its birth to its death.

**M**^3^: Multi-CAM metaFuME Markov (model). A model based on the metaFuME set *𝒟*_Λ_ of CAM-specific distributions (Eq. 10) and CAM-specific Markov transition matrices *P*_*λ*_, *λ* = 1, …, Λ, derived from the set 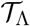 of metaFuME sequence index set ensembles (see Fig. 3). This model contains only local movement information and cannot be used to simulate movement patterns motivated by headings to distant targets.

**MetaFuME:** A correlated stereotypical or characteristic sequence of FuMEs of fixed duration equal to the time between consecutive relocation points (only applicable to relatively high frequency data: ideally, metaFuMEs should contain no more than several tens of FuMES).

**PS:** Path Segmentation. This is the process of breaking up a movement path into reliably stable metaFuMEs, CAMs, DARs, and LiMPs using a suite of methods that include BCPA and HMM approaches (Nams 2014, Edelhoff et al. 2016, Seidel et al. 2018).

**SL:** Step Length. The distance between consecutive relocation points, as generated from Eq. 1 using Eq. 3.

**TA:** Turning Angle. The change in the angle of heading across three consecutive relocation points, as generated from Eq. 1 using Eq. 5.

## Acknowledgements

I thank Briana Abrahms, Shirli Bar-David, Paul Cross, Victoria Goodall, Nir Horvitz, Leif Loe, Ran Nathan, Norman Owen-Smith, Leo Polansky, David Saltz, Dana Seidel, Orr Spiegel, Krti Tallam, Miriam Tsalyuk, George Wittemyer for comments on earlier versions of this paper. All inaccuracies and misconceptions are my own.

## Notes

#### Summary of Updates

First revision: Add a second panel to Fig. 4, minor changes to Fig. 1, and minor edits to the table and text. Change the in-text citation style. Second revision: Eq 13 corrected. New figure added (Fig 4; old Fig. 4 now Fig. 5). Provided clarification on simulation models and named it Markov metaFuME movement model. Third revision: update discussion and streamlined text including discussion of M-cubed (multi-CAM, metaFuME Markov) model. Four revision: Added definition of metaFuME to glossary

## References

Abrahms, B., Seidel, D. P., Dougherty, E., Hazen, E. L., Bograd, S. J., Wilson, A. M., McNutt, J. W., Costa, D. P., Blake, S., Brashares, J. S. et al. (2017), ‘Suite of simple metrics reveals common movement syndromes across vertebrate taxa’, Movement ecology 5(1), 12.

Allen, R. M., Metaxas, A. & Snelgrove, P. V. (2018), ‘Applying movement ecology to marine animals with complex life cycles’, Annual Review of Marine Science 10, 19–42.

Andersen, T., Carstensen, J., Hernandez-Garcia, E. & Duarte, C. M. (2009), ‘Ecological thresholds and regime shifts: approaches to identification’, Trends in Ecology & Evolution 24(1), 49–57.

Avgar, T., Potts, J. R., Lewis, M. A. & Boyce, M. S. (2016), ‘Integrated step selection analysis: bridging the gap between resource selection and animal movement’, Methods in Ecology and Evolution 7(5), 619–630.

Bar-David, S., Bar-David, I., Cross, P. C., Ryan, S. J., Knechtel, C. U. & Getz, W. M. (2009), ‘Methods for assessing movement path recursion with application to African buffalo in South Africa’, Ecology 90(9), 2467–2479.

Bartumeus, F., da Luz, M. G. E., Viswanathan, G. M. & Catalan, J. (2005), ‘Animal search strategies: a quantitative random-walk analysis’, Ecology 86(11), 3078–3087.

Bennison, A., Bearhop, S., Bodey, T. W., Votier, S. C., Grecian, W. J., Wakefield, E. D., Hamer, K. C. & Jessopp, M. (2018), ‘Search and foraging behaviors from movement data: a comparison of methods’, Ecology and evolution 8(1), 13–24.

Berger-Tal, O. & Bar-David, S. (2015), ‘Recursive movement patterns: review and synthesis across species’, Ecosphere 6(9), 1–12.

Bergman, C. M., Schaefer, J. A. & Luttich, S. (2000), ‘Caribou movement as a correlated random walk’, Oecologia 123(3), 364–374.

Bischof, R., Loe, L. E., Meisingset, E. L., Zimmermann, B., Van Moorter, B. & Mysterud, A. (2012), ‘A migratory northern ungulate in the pursuit of spring: jumping or surfing the green wave?’, The American Naturalist 180(4), 407–424.

Bork, E. W. & Su, J. G. (2007), ‘Integrating lidar data and multispectral imagery for enhanced classification of rangeland vegetation: A meta analysis’, Remote Sensing of Environment 111(1), 11–24.

Bowler, D. E. & Benton, T. G. (2005), ‘Causes and consequences of animal dispersal strategies: relating individual behaviour to spatial dynamics’, Biological Reviews 80(2), 205–225.

Boyce, M. S., Vernier, P. R., Nielsen, S. E. & Schmiegelow, F. K. (2002), ‘Evaluating resource selection functions’, Ecological modelling 157(2-3), 281–300.

Brodie, J. F., Giordano, A. J., Dickson, B., Hebblewhite, M., Bernard, H., Mohd-Azlan, J., Anderson, J. & Ambu, L. (2015), ‘Evaluating multispecies landscape connectivity in a threatened tropical mammal community’, Conservation Biology 29(1), 122–132.

Bunnefeld, N., Börger, L., van Moorter, B., Rolandsen, C. M., Dettki, H., Solberg, E. J. & Ericsson, G. (2011), ‘A model-driven approach to quantify migration patterns: individual, regional and yearly differences’, Journal of Animal Ecology 80(2), 466–476.

Byers, J. A. (2001), ‘Correlated random walk equations of animal dispersal resolved by simulation’, Ecology 82(6), 1680–1690.

Calabrese, J. M., Fleming, C. H. & Gurarie, E. (2016), ‘ctmm: an r package for analyzing animal relocation data as a continuous-time stochastic process’, Methods in Ecology and Evolution 7(9), 1124–1132.

Cattarino, L., McAlpine, C. A. & Rhodes, J. R. (2016), ‘Spatial scale and movement behaviour traits control the impacts of habitat fragmentation on individual fitness’, Journal of Animal Ecology 85(1), 168–177.

Chakravarty, P., Maalberg, M., Cozzi, G., Ozgul, A. & Aminian, K. (2019), ‘Behavioural compass: animal behaviour recognition using magnetometers’, Movement Ecology 7(1), 28.

Chatfield, C. (2016), The analysis of time series: an introduction, Chapman and Hall/CRC.

Chen, J. & Gupta, A. K. (2011), Parametric statistical change point analysis: with applications to genetics, medicine, and finance, Springer Science & Business Media.

Coello, C. A. C. (2003), Evolutionary multi-objective optimization: A critical review, in ‘Evolutionary optimization’, Springer, pp. 117–146.

Cross, P. C., Almberg, E. S., Haase, C. G., Hudson, P. J., Maloney, S. K., Metz, M. C., Munn, A. J., Nugent, P., Putzeys, O., Stahler, D. R. et al. (2016), ‘Energetic costs of mange in wolves estimated from infrared thermography’, Ecology 97(8), 1938–1948.

Cross, P. C., Edwards, W. H., Scurlock, B. M., Maichak, E. J. & Rogerson, J. D. (2007), ‘Effects of management and climate on elk brucellosis in the greater yellowstone ecosystem’, Ecological Applications 17(4), 957–964.

de Weerd, N., van Langevelde, F., van Oeveren, H., Nolet, B. A., Kölzsch, A., Prins, H. H. & de Boer, W. F. (2015), ‘Deriving animal behaviour from high-frequency gps: tracking cows in open and forested habitat’, Plos one 10(6), e0129030.

DeCesare, N. J., Hebblewhite, M., Schmiegelow, F., Hervieux, D., McDermid, G. J., Neufeld, L., Bradley, M., Whittington, J., Smith, K. G., Morgantini, L. E. et al. (2012), ‘Transcending scale dependence in identifying habitat with resource selection functions’, Ecological Applications 22(4), 1068–1083.

Delcourt, J., Denöel, M., Ylieff, M. & Poncin, P. (2013), ‘Video multitracking of fish behaviour: a synthesis and future perspectives’, Fish and Fisheries 14(2), 186–204.

Delp, S. L. & Loan, J. P. (2000), ‘A computational framework for simulating and analyzing human and animal movement’, Computing in Science & Engineering 2(5), 46–55.

Donati, G., Campera, M., Balestri, M., Serra, V., Barresi, M., Schwitzer, C., Curtis, D. J. & Santini, L. (2016), ‘Ecological and anthropogenic correlates of activity patterns in eulemur’, International Journal of Primatology 37(1), 29–46.

Dougherty, E. R., Seidel, D. P., Carlson, C. J., Spiegel, O. & Getz, W. M. (2018), ‘Going through the motions: incorporating movement analyses into disease research’, Ecology letters 21(4), 588–604.

Dovera, L. & Della Rossa, E. (2011), ‘Multimodal ensemble kalman filtering using gaussian mixture models’, Computational Geosciences 15(2), 307–323.

Edelhoff, H., Signer, J. & Balkenhol, N. (2016), ‘Path segmentation for beginners: an overview of current methods for detecting changes in animal movement patterns’, Movement ecology 4(1), 21.

Fahr, J., Abedi-Lartey, M., Esch, T., Machwitz, M., Suu-Ire, R., Wikelski, M. & Dechmann, D. K. (2015), ‘Pronounced seasonal changes in the movement ecology of a highly gregarious central-place forager, the african straw-coloured fruit bat (eidolon helvum)’, PloS one 10(10), e0138985.

Fauchald, P. & Tveraa, T. (2003), ‘Using first-passage time in the analysis of area-restricted search and habitat selection’, Ecology 84(2), 282–288.

Fehlmann, G., ORiain, M. J., Hopkins, P. W., OSullivan, J., Holton, M. D., Shepard, E. L. & King, A. J. (2017), ‘Identification of behaviours from accelerometer data in a wild social primate’, Animal Biotelemetry 5(1), 6.

Fleming, C. H. & Calabrese, J. M. (2017), ‘A new kernel density estimator for accurate home-range and species-range area estimation’, Methods in Ecology and Evolution 8(5), 571–579.

Fleming, C. H., Calabrese, J. M., Mueller, T., Olson, K. A., Leimgruber, P. & Fagan, W. F. (2014), ‘From fine-scale foraging to home ranges: a semivariance approach to identifying movement modes across spatiotemporal scales’, The American Naturalist 183(5), E154–E167.

Fortin, D., Beyer, H. L., Boyce, M. S., Smith, D. W., Duchesne, T. & Mao, J. S. (2005), ‘Wolves influence elk movements: behavior shapes a trophic cascade in yellowstone national park’, Ecology 86(5), 1320–1330.

Fortin, D., Boyce, M. S., Merrill, E. H. & Fryxell, J. M. (2004), ‘Foraging costs of vigilance in large mammalian herbivores’, Oikos 107(1), 172–180.

Franke, A., Caelli, T. & Hudson, R. J. (2004), ‘Analysis of movements and behavior of caribou (rangifer tarandus) using hidden markov models’, Ecological Modelling 173(2-3), 259–270.

Getz, W. M., Marshall, C. R., Carlson, C. J., Giuggioli, L., Ryan, S. J., Romañach, S. S., Boettiger, C., Chamberlain, S. D., Larsen, L., DOdorico, P. et al. (2018), ‘Making ecological models adequate’, Ecology letters 21(2), 153–166.

Getz, W. M., Salter, R. & Tallam, K. (2019), ‘A quantitative narrative on movement, disease and patch exploitation in nesting agent groups’, bioRxiv. URL: https://www.biorxiv.org/content/early/2019/10/03/791400

Getz, W. M. & Saltz, D. (2008), ‘A framework for generating and analyzing movement paths on ecological landscapes’, Proceedings of the National Academy of Sciences 105(49), 19066–19071.

Getz, W., Vissat, L. L. & Salter, R. (2019), ‘Simulation and analysis of animal movement paths using numerus model builder’, bioRxiv. URL: https://www.biorxiv.org/content/early/2019/12/15/2019.12.15.876987

Gevers, D., Cohan, F. M., Lawrence, J. G., Spratt, B. G., Coenye, T., Feil, E. J., Stackebrandt, E., Van de Peer, Y., Vandamme, P., Thompson, F. L. et al. (2005), ‘Re-evaluating prokaryotic species’, Nature Reviews Microbiology 3(9), 733.

Giotto, N., Gerard, J.-F., Ziv, A., Bouskila, A. & Bar-David, S. (2015), ‘Space-use patterns of the asiatic wild ass (*equus hemionus*): complementary insights from displacement, recursion movement and habitat selection analyses’, PloS one 10(12), e0143279.

Goodall, V., Fatti, L. & Owen-Smith, N. (2017), ‘Animal movement modelling: independent or dependent models?’, South African Statistical Journal 51(2), 295–315.

Grémillet, D. & Boulinier, T. (2009), ‘Spatial ecology and conservation of seabirds facing global climate change: a review’, Marine Ecology Progress Series 391, 121–137.

Gurarie, E., Andrews, R. D. & Laidre, K. L. (2009), ‘A novel method for identifying behavioural changes in animal movement data’, Ecology letters 12(5), 395–408.

Gurarie, E., Bracis, C., Delgado, M., Meckley, T. D., Kojola, I. & Wagner, C. M. (2016), ‘What is the animal doing? tools for exploring behavioural structure in animal movements’, Journal of Animal Ecology 85(1), 69–84.

Gurarie, E., Fleming, C. H., Fagan, W. F., Laidre, K. L., Hernáandez-Pliego, J. & Ovaskainen, O. (2017), ‘Correlated velocity models as a fundamental unit of animal movement: synthesis and applications’, Movement ecology 5(1), 13.

Hardin, P. E. & Panda, S. (2013), ‘Circadian timekeeping and output mechanisms in animals’, Current opinion in neurobiology 23(5), 724–731.

Harel, R., Duriez, O., Spiegel, O., Fluhr, J., Horvitz, N., Getz, W. M., Bouten, W., Sarrazin, F., Hatzofe, O. & Nathan, R. (2016), ‘Decision-making by a soaring bird: time, energy and risk considerations at different spatio-temporal scales’, Philosophical Transactions of the Royal Society B: Biological Sciences 371(1704), 20150397.

Harel, R., Horvitz, N. & Nathan, R. (2016), ‘Adult vultures outperform juveniles in challenging thermal soaring conditions’, Scientific reports 6, 27865.

Harel, R. & Nathan, R. (2018), ‘The characteristic time-scale of perceived information for decision-making: Departure from thermal columns in soaring birds’, Functional ecology 32(8), 2065–2072.

Harel, R., Spiegel, O., Getz, W. M. & Nathan, R. (2017), ‘Social foraging and individual consistency in following behaviour: testing the information centre hypothesis in free-ranging vultures’, Proceedings of the Royal Society B: Biological Sciences 284(1852), 20162654.

Hays, G. C., Ferreira, L. C., Sequeira, A. M., Meekan, M. G., Duarte, C. M., Bailey, H., Bailleul, F., Bowen, W. D., Caley, M. J., Costa, D. P. et al. (2016), ‘Key questions in marine megafauna movement ecology’, Trends in ecology & evolution 31(6), 463–475.

Hein, A. M., Carrara, F., Brumley, D. R., Stocker, R. & Levin, S. A. (2016), ‘Natural search algorithms as a bridge between organisms, evolution, and ecology’, Proceedings of the National Academy of Sciences 113(34), 9413–9420.

Hinton, G., Deng, L., Yu, D., Dahl, G. E., Mohamed, A.-r., Jaitly, N., Senior, A., Vanhoucke, V., Nguyen, P., Sainath, T. N. et al. (2012), ‘Deep neural networks for acoustic modeling in speech recognition: The shared views of four research groups’, IEEE Signal processing magazine 29(6), 82–97.

Hurme, E., Gurarie, E., Greif, S., Flores-Martínez, J. J., Wilkinson, G. S., Yovel, Y. et al. (2019), ‘Acoustic evaluation of behavioral states predicted from gps tracking: a case study of a marine fishing bat’, Movement ecology 7(1), 21.

Johnson, C. J., Parker, K. L., Heard, D. C. & Gillingham, M. P. (2002), ‘A multiscale behavioral approach to understanding the movements of woodland caribou’, Ecological Applications 12(6), 1840–1860.

Johnson, D. S., London, J. M., Lea, M.-A. & Durban, J. W. (2008), ‘Continuous-time correlated random walk model for animal telemetry data’, Ecology 89(5), 1208–1215.

Jonsen, I. D., Myers, R. A. & Flemming, J. M. (2003), ‘Meta-analysis of animal movement using state-space models’, Ecology 84(11), 3055–3063.

Jonsen, I. D., Myers, R. A. & James, M. C. (2006), ‘Robust hierarchical state–space models reveal diel variation in travel rates of migrating leatherback turtles’, Journal of Animal Ecology 75(5), 1046–1057.

Jonsen, I. D., Myers, R. A. & James, M. C. (2007), ‘Identifying leatherback turtle foraging behaviour from satellite telemetry using a switching state-space model’, Marine Ecology Progress Series 337, 255–264.

Kane, A. S., Salierno, J. D., Gipson, G. T., Molteno, T. C. & Hunter, C. (2004), ‘A video-based movement analysis system to quantify behavioral stress responses of fish’, Water Research 38(18), 3993–4001.

Kareiva, P. & Shigesada, N. (1983), ‘Analyzing insect movement as a correlated random walk’, Oecologia 56(2-3), 234–238.

Killick, R. & Eckley, I. (2014), ‘changepoint: An r package for changepoint analysis’, Journal of statistical software 58(3), 1–19.

Langrock, R., King, R., Matthiopoulos, J., Thomas, L., Fortin, D. & Morales, J. M. (2012), ‘Flexible and practical modeling of animal telemetry data: hidden markov models and extensions’, Ecology 93(11), 2336–2342.

Larsen, L. G., Eppinga, M. B., Passalacqua, P., Getz, W. M., Rose, K. A. & Liang, M. (2016), ‘Appropriate complexity landscape modeling’, Earth-science reviews 160, 111–130.

Levin, S. A. (1992), ‘The problem of pattern and scale in ecology: the robert h. macarthur award lecture’, Ecology 73(6), 1943–1967.

Lyons, A. J., Turner, W. C. & Getz, W. M. (2013), ‘Home range plus: a space-time characterization of movement over real landscapes’, Movement Ecology 1(1), 2.

Marin, J.-M., Pudlo, P., Robert, C. P. & Ryder, R. J. (2012), ‘Approximate bayesian computational methods’, Statistics and Computing 22(6), 1167–1180.

Marra, P. P., Cohen, E. B., Loss, S. R., Rutter, J. E. & Tonra, C. M. (2015), ‘A call for full annual cycle research in animal ecology’, Biology letters 11(8), 20150552.

Matteson, D. S. & James, N. A. (2014), ‘A nonparametric approach for multiple change point analysis of multivariate data’, Journal of the American Statistical Association 109(505), 334–345.

McCulloch, C. & Cain, M. (1989), ‘Analyzing discrete movement data as a correlated random walk’, Ecology 70(2), 383–388.

McGarigal, K., Wan, H. Y., Zeller, K. A., Timm, B. C. & Cushman, S. A. (2016), ‘Multi-scale habitat selection modeling: a review and outlook’, Landscape ecology 31(6), 1161–1175.

McGavin, S. L., Bishop-Hurley, G. J., Charmley, E., Greenwood, P. L. & Callaghan, M. J. (2018), ‘Effect of gps sample interval and paddock size on estimates of distance travelled by grazing cattle in rangeland, australia’, The Rangeland Journal 40(1), 55–64.

McKenzie, H. W., Lewis, M. A. & Merrill, E. H. (2009), ‘First passage time analysis of animal movement and insights into the functional response’, Bulletin of mathematical biology 71(1), 107–129.

Michelot, T., Langrock, R. & Patterson, T. A. (2016), ‘movehmm: An r package for the statistical modelling of animal movement data using hidden markov models’, Methods in Ecology and Evolution 7(11), 1308–1315.

Milner-Gulland, E., Fryxell, J. M. & Sinclair, A. R. (2011), Animal migration: a synthesis, Oxford University Press.

Moore, E. R., Mihaylova, S. A., Vandamme, P., Krichevsky, M. I. & Dijkshoorn, L. (2010), ‘Microbial systematics and taxonomy: relevance for a microbial commons’, Research in microbiology 161(6), 430–438.

Morales, J. M., Haydon, D. T., Frair, J., Holsinger, K. E. & Fryxell, J. M. (2004), ‘Extracting more out of relocation data: building movement models as mixtures of random walks’, Ecology 85(9), 2436–2445.

Morelle, K., Podgórski, T., Prévot, C., Keuling, O., Lehaire, F. & Lejeune, P. (2015), ‘Towards understanding wild boar *sus scrofa* movement: a synthetic movement ecology approach’, Mammal Review 45(1), 15–29.

Morellet, N., Bonenfant, C., Börger, L., Ossi, F., Cagnacci, F., Heurich, M., Kjellander, P., Linnell, J. D., Nicoloso, S., Sustr, P. et al. (2013), ‘Seasonality, weather and climate affect home range size in roe deer across a wide latitudinal gradient within e urope’, Journal of Animal Ecology 82(6), 1326–1339.

Mueller, T. & Fagan, W. F. (2008), ‘Search and navigation in dynamic environments–from individual behaviors to population distributions’, Oikos 117(5), 654–664.

Myers, J. H. (2018), ‘Population cycles: generalities, exceptions and remaining mysteries’, Proceedings of the Royal Society B: Biological Sciences 285(1875), 20172841.

Mysterud, A., Stenseth, N. C., Yoccoz, N. G., Langvatn, R. & Steinheim, G. (2001), ‘Nonlinear effects of large-scale climatic variability on wild and domestic herbivores’, Nature 410(6832), 1096.

Nams, V. O. (2014), ‘Combining animal movements and behavioural data to detect behavioural states’, Ecology letters 17(10), 1228–1237.

Nathan, R., Getz, W. M., Revilla, E., Holyoak, M., Kadmon, R., Saltz, D. & Smouse, P. E. (2008), ‘A movement ecology paradigm for unifying organismal movement research’, Proceedings of the National Academy of Sciences 105(49), 19052–19059.

Nathan, R., Spiegel, O., Fortmann-Roe, S., Harel, R., Wikelski, M. & Getz, W. M. (2012), ‘Using triaxial acceleration data to identify behavioral modes of free-ranging animals: general concepts and tools illustrated for griffon vultures’, Journal of Experimental Biology 215(6), 986–996.

Northrup, J. M., Avrin, A., Anderson, C. R., Brown, E. & Wittemyer, G. (2019), ‘On-animal acoustic monitoring provides insight to ungulate foraging behavior’, Journal of Mammalogy.

Odu, G. & Charles-Owaba, O. (2013), ‘Review of multi-criteria optimization methods–theory and applications’, IOSR Journal of Engineering (IOSRJEN) 3(10), 1–14.

Owen-Smith, N. (2013), ‘Daily movement responses by african savanna ungulates as an indicator of seasonal and annual food stress’, Wildlife Research 40(3), 232–240.

Owen-Smith, N., Fryxell, J. & Merrill, E. (2010), ‘Foraging theory upscaled: the behavioural ecology of herbivore movement’, Philosophical Transactions of the Royal Society B: Biological Sciences 365(1550), 2267–2278.

Owen-Smith, N. & Goodall, V. (2014), ‘Coping with savanna seasonality: comparative daily activity patterns of a frican ungulates as revealed by gps telemetry’, Journal of Zoology 293(3), 181–191.

Owen-Smith, N. & Goodall, V. (2019), ‘Movement informatics of large mammalian herbivores across hierarchical scales: inferring biology from statistics’, Under review.

Owen-Smith, N. & Martin, J. (2015), ‘Identifying space use at foraging arena scale within the home ranges of large herbivores’, PloS one 10(6), e0128821.

Panzacchi, M., Van Moorter, B., Strand, O., Saerens, M., Kivimäki, I., St. Clair, C. C., Herfindal, I. & Boitani, L. (2016), ‘Predicting the continuum between corridors and barriers to animal movements using step selection functions and randomized shortest paths’, Journal of Animal Ecology 85(1), 32–42.

Patterson, T. A., Thomas, L., Wilcox, C., Ovaskainen, O. & Matthiopoulos, J. (2008), ‘State–space models of individual animal movement’, Trends in ecology & evolution 23(2), 87–94.

Pettorelli, N., Laurance, W. F., O’Brien, T. G., Wegmann, M., Nagendra, H. & Turner, W. (2014), ‘Satellite remote sensing for applied ecologists: opportunities and challenges’, Journal of Applied Ecology 51(4), 839–848.

Pettorelli, N., Ryan, S., Mueller, T., Bunnefeld, N., Jedrzejewska, B., Lima, M. & Kausrud, K. (2011), ‘The normalized difference vegetation index (ndvi): unforeseen successes in animal ecology’, Climate Research 46(1), 15–27.

Pohle, J., Langrock, R., van Beest, F. M. & Schmidt, N. M. (2017), ‘Selecting the number of states in hidden markov models: pragmatic solutions illustrated using animal movement’, Journal of Agricultural, Biological and Environmental Statistics 22(3), 270–293.

Polansky, L., Kilian, W. & Wittemyer, G. (2015), ‘Elucidating the significance of spatial memory on movement decisions by african savannah elephants using state–space models’, Proceedings of the Royal Society B: Biological Sciences 282(1805), 20143042.

Polansky, L., Wittemyer, G., Cross, P. C., Tambling, C. J. & Getz, W. M. (2010), ‘From moonlight to movement and synchronized randomness: Fourier and wavelet analyses of animal location time series data’, Ecology 91(5), 1506–1518.

Preisler, H. K., Ager, A. A., Johnson, B. K. & Kie, J. G. (2004), ‘Modeling animal movements using stochastic differential equations’, Environmetrics 15(7), 643–657.

Rahimi, S. & Owen-Smith, N. (2007), ‘Movement patterns of sable antelope in the kruger national park from gps/gsm collars: a preliminary assessment’, African Journal of Wildlife Research 37(2), 143–152.

Rosselló-Móra, R. & Whitman, W. B. (2019), ‘Dialogue on the nomenclature and classification of prokaryotes’, Systematic and applied microbiology 42(1), 5–14.

Runge, C. A., Martin, T. G., Possingham, H. P., Willis, S. G. & Fuller, R. A. (2014), ‘Conserving mobile species’, Frontiers in Ecology and the Environment 12(7), 395–402.

Sapir, N., Wikelski, M., McCue, M. D., Pinshow, B. & Nathan, R. (2010), ‘Flight modes in migrating european bee-eaters: heart rate may indicate low metabolic rate during soaring and gliding’, PLoS One 5(11), e13956.

Schuh, R. T. (2003), ‘The linnaean system and its 250-year persistence’, The Botanical Review 69(1), 59.

Seebacher, F. & Post, E. (2015), ‘Climate change impacts on animal migration’, Climate Change Responses 2(1), 5.

Seidel, D. P., Dougherty, E., Carlson, C. & Getz, W. M. (2018), ‘Ecological metrics and methods for gps movement data’, International Journal of Geographical Information Science 32(11), 2272–2293.

Shepard, E. L., Wilson, R. P., Rees, W. G., Grundy, E., Lambertucci, S. A. & Vosper, S. B. (2013), ‘Energy landscapes shape animal movement ecology’, The American Naturalist 182(3), 298–312.

Siniff, D. B. & Jessen, C. (1969), A simulation model of animal movement patterns, in ‘Advances in ecological research’, Vol. 6, Elsevier, pp. 185–219.

Spiegel, O., Harel, R., Centeno-Cuadros, A., Hatzofe, O., Getz, W. M. & Nathan, R. (2015), ‘Moving beyond curve fitting: using complementary data to assess alternative explanations for long movements of three vulture species’, The American Naturalist 185(2), E44–E54.

Spiegel, O. & O’Farrell, S. (2019), ‘Spatial orientation and time: Methods’, Encyclopedia of Animal Behavior pp. 518–528.

Spink, A., Tegelenbosch, R., Buma, M. & Noldus, L. (2001), ‘The ethovision video tracking systema tool for behavioral phenotyping of transgenic mice’, Physiology & behavior 73(5), 731–744.

Symonds, M. R. & Moussalli, A. (2011), ‘A brief guide to model selection, multimodel inference and model averaging in behavioural ecology using akaikes information criterion’, Behavioral Ecology and Sociobiology 65(1), 13–21.

Takahashi, J. S., Hong, H.-K., Ko, C. H. & McDearmon, E. L. (2008), ‘The genetics of mammalian circadian order and disorder: implications for physiology and disease’, Nature reviews genetics 9(10), 764.

Thessen, A. (2016), ‘Adoption of machine learning techniques in ecology and earth science’, One Ecosystem 1, e8621.

Thurfjell, H., Ciuti, S. & Boyce, M. S. (2014), ‘Applications of step-selection functions in ecology and conservation’, Movement ecology 2(1), 4.

Toledo, S., Kishon, O., Orchan, Y., Shohat, A. & Nathan, R. (2016), Lessons and experiences from the design, implementation, and deployment of a wildlife tracking system, in ‘Software Science, Technology and Engineering (SWSTE), 2016 IEEE International Conference on’, IEEE, pp. 51–60.

Torney, C. J., Hopcraft, J. G. C., Morrison, T. A., Couzin, I. D. & Levin, S. A. (2018), ‘From single steps to mass migration: the problem of scale in the movement ecology of the serengeti wildebeest’, Philosophical Transactions of the Royal Society B: Biological Sciences 373(1746), 20170012.

Torres, L. G., Orben, R. A., Tolkova, I. & Thompson, D. R. (2017), ‘Classification of animal movement behavior through residence in space and time’, PloS one 12(1), e0168513.

Townsend Peterson, A., Papeş, M. & Eaton, M. (2007), ‘Transferability and model evaluation in ecological niche modeling: a comparison of garp and maxent’, Ecography 30(4), 550–560.

Tracey, J. A., Bevins, S. N., VandeWoude, S. & Crooks, K. R. (2014), ‘An agent-based movement model to assess the impact of landscape fragmentation on disease transmission’, Ecosphere 5(9), 1–24.

Tsalyuk, M., Kilian, W., Reineking, B. & Getz, W. M. (2019), ‘Temporal variation in resource selection of african elephants follows long-term variability in resource availability’, Ecological Monographs 89(2), e01348.

Turchin, P. (1998), Quantitative analysis of movement, Sinauer assoc. Sunderland (mass.).

Valletta, J. J., Torney, C., Kings, M., Thornton, A. & Madden, J. (2017), ‘Applications of machine learning in animal behaviour studies’, Animal Behaviour 124, 203–220.

Van Moorter, B., Visscher, D. R., Jerde, C. L., Frair, J. L. & Merrill, E. H. (2010), ‘Identifying movement states from location data using cluster analysis’, The Journal of Wildlife Management 74(3), 588–594.

Vissat, L. L., Salter, R. & Getz, W. M. (2019), ‘Simulation and analysis of animal movement paths using numerus model builder’, bioRxiv. URL: https://www.biorxiv.org/content/early/2019/xxxx

Walther, G. et al. (2009), ‘Inference and modeling with log-concave distributions’, Statistical Science 24(3), 319–327.

Wang, Y., Nickel, B., Rutishauser, M., Bryce, C. M., Williams, T. M., Elkaim, G. & Wilmers, C. C. (2015), ‘Movement, resting, and attack behaviors of wild pumas are revealed by tri-axial accelerometer measurements’, Movement Ecology 3(1), 2.

Weiser, A. W., Orchan, Y., Nathan, R., Charter, M., Weiss, A. J. & Toledo, S. (2016), Characterizing the accuracy of a self-synchronized reverse-gps wildlife localization system, in ‘Information Processing in Sensor Networks (IPSN), 2016 15th ACM/IEEE International Conference on’, IEEE, pp. 1–12.

Williams, H. J., Holton, M. D., Shepard, E. L., Largey, N., Norman, B., Ryan, P. G., Duriez, O., Scantlebury, M., Quintana, F., Magowan, E. A. et al. (2017), ‘Identification of animal movement patterns using tri-axial magnetometry’, Movement ecology 5(1), 6.

Williams, H. J., Taylor, L. A., Benhamou, S., Bijleveld, A. I., Clay, T. A., de Grissac, S., Demšar, U., English, H. M., Franconi, N., Gómez-Laich, A. et al. (2019), ‘Optimising the use of bio-loggers for movement ecology research’, Journal of Animal Ecology.

Wilmers, C. C., Nickel, B., Bryce, C. M., Smith, J. A., Wheat, R. E. & Yovovich, V. (2015), ‘The golden age of bio-logging: how animal-borne sensors are advancing the frontiers of ecology’, Ecology 96(7), 1741–1753.

Wittemyer, G., Northrup, J. M. & Bastille-Rousseau, G. (2019), ‘Behavioural valuation of landscapes using movement data’, Philosophical Transactions of the Royal Society B 374(1781), 20180046.

Wittemyer, G., Polansky, L., Douglas-Hamilton, I. & Getz, W. M. (2008), ‘Disentangling the effects of forage, social rank, and risk on movement autocorrelation of elephants using fourier and wavelet analyses’, Proceedings of the National Academy of Sciences pp. pnas–0801744105.

Worton, B. (1987), ‘A review of models of home range for animal movement’, Ecological modelling 38(3-4), 277–298.

Yerushalmi, S. & Green, R. M. (2009), ‘Evidence for the adaptive significance of circadian rhythms’, Ecology letters 12(9), 970–981.

Zeller, K. A., McGarigal, K. & Whiteley, A. R. (2012), ‘Estimating landscape resistance to movement: a review’, Landscape ecology 27(6), 777–797.

Zhang, J., Hull, V., Ouyang, Z., He, L., Connor, T., Yang, H., Huang, J., Zhou, S., Zhang, Z., Zhou, C. et al. (2017), ‘Modeling activity patterns of wildlife using time-series analysis’, Ecology and evolution 7(8), 2575–2584.

Zhang, Z., Geiger, J., Pohjalainen, J., Mousa, A. E.-D., Jin, W. & Schuller, B. (2018), ‘Deep learning for environmentally robust speech recognition: An overview of recent developments’, ACM Transactions on Intelligent Systems and Technology (TIST) 9(5), 49.

Zhao, K. & Jurdak, R. (2016), ‘Understanding the spatiotemporal pattern of grazing cattle movement’, Scientific reports 6, 31967.

Zidon, R., Garti, S., Getz, W. M. & Saltz, D. (2017), ‘Zebra migration strategies and anthrax in etosha national park, namibia’, Ecosphere 8(8), e01925.

Zucchini, W., MacDonald, I. L. & Langrock, R. (2016), Hidden Markov models for time series: an introduction using R, Chapman and Hall/CRC.

